# EPEPDI: prediction of binding free energy changes from missense mutations in double and single-stranded DNA-binding proteins

**DOI:** 10.1101/2025.07.08.663633

**Authors:** Xuan Yu

## Abstract

Predicting changes in binding free energy due to missense mutations (MMs) in protein-DNA interactions (PDIs) is vital for understanding disease mechanisms and advancing therapeutic strategies. However, many existing models fail to account for the unique characteristics of MMs in double-stranded DNA binding proteins (DSBs) and single-stranded DNA binding proteins (SSBs). To address this, we constructed a comprehensive dataset from diverse sources, clearly distinguishing between DSBs and SSBs. Using sequence-based embeddings from pre-trained protein language models, including ESM2, ProtTrans, and ESM1v, we developed EPEPDI, a deep learning framework that integrates these embeddings through a multi-channel architecture. To refine predictive accuracy, we introduced an information entropy-based algorithm, determining 181 residues as the optimal sequence length where amino acid contributions and entropy dynamics balance. This approach boosts both precision and computational efficiency, enabling scalable analysis of mutation impacts on DNA-binding proteins. Ablation studies validated optimal feature combinations, demonstrating that EPEPDI outperforms existing approaches, achieving an average Pearson correlation coefficient of 0.755 on the MPD276 dataset via ten-fold cross-validation and 0.632 on independent tests for both DSBs and SSBs. This work highlights the importance of distinguishing DSBs and SSBs in PDIs and shows the potential of advanced machine learning in biological research.

## 1. Introduction

Protein-DNA interactions (PDIs) are essential for key cellular processes such as DNA replication, repair, and recombination [1–3]. Accurately predicting binding affinities is essential to unravel biochemical mechanisms, including drug efficacy [4], antibody-antigen interactions [5], and early disease detection [6]. These affinities depend on intermolecular forces (e.g., electrostatics, van der Waals forces, hydrogen bonding, and hydrophobic effects) and protein structural features [7–10]. Missense mutations in these DNA-binding proteins can affect these determinants, leading to significant alterations in binding free energies [11]. Laboratory techniques like isothermal titration calorimetry, electrophoretic mobility shift assay, and systematic evolution of ligands by exponential enrichment offer precise measurements of binding free energy changes (ΔΔG) [12–14]. However, these methods are costly, time-intensive, and impractical for large-scale genomic data analysis. Consequently, computational approaches have gained prominence for efficient ΔΔG prediction. Among existing methods, FlodX utilizes high-resolution 3D structures to model and predict the impact of mutations on binding energies, which is crucial for understanding mutation-induced ΔΔG changes [10, 15, 16].

Recent research has advanced our understanding of protein-nucleotide interactions, including PDIs and protein-RNA interactions (PRIs). For instance, ProNIT provides thermodynamic data such as free energy and dissociation constants [17]. Building on this, SAMPDI employs a modified molecular mechanics Poisson Boltzmann surface area method to predict ΔΔG for MPDs [11], while PremPDI uses molecular mechanics and rapid side-chain optimization, trained via multiple linear regression, to assess mutation impacts and model mutant structures [18]. Beyond PDIs, studies have also examined mutation effects on PRIs. Tools like mCSM-NA integrate pharmacophore modeling and nucleic acid properties for prediction [19], which PEMPNI conducts systematic comparisons between mutations in DNA-binding proteins (MPDs) and RNA-binding proteins (MPRs), using specialized regression techniques and geometric energy features [20]. Together, these efforts establish a robust framework for analyzing how missense mutations affect protein-nucleotide interactions, offering insights into their structural and functional consequences.

While significant progress has been made in understanding protein-DNA interactions (PDIs), a critical distinction is often overlooked: DNA-binding proteins encompass both double-stranded DNA binding proteins (DSBs) and single-stranded DNA binding proteins (SSBs) [21–23]. These two categories exhibit marked differences in their functions, DNA structure preference, and binding mechanisms. Previous studies highlight that DSBs and SSBs differ not only in sequence but also in their structural characteristics, leading to the development of classifiers tailored to these distinctions [24, 25]. SSBs are essential for DNA replication, repair, and recombination, protecting single-stranded DNA and preventing secondary structure formation [26–28]. They bind with high affinity and can reposition along the strand [29]. In contrast, DSBs typically act as transcription factors or DNA repair proteins, maintaining genomic stability [30]. However, existing models for predicting the effects of missense mutations on PDIs frequently utilize mixed datasets from both DSBs and SSBs, potentially reducing accuracy. Moreover, these models depend on energy-based and structural features that require precise structural data and often fail to capture subtle molecular interactions [11, 20]. Recent advances in large-scale language models provide atomic-resolution protein representations, such as ESM features [31], offering a promising alternative for improving model prediction ability.

Recognizing the distinct binding mechanisms of DSBs and SSBs, we separately analyze the effects of missense mutations on these two categories to enhance the prediction of ΔΔG changes in PDIs. In this study, we constructed a comprehensive dataset (distinguish between DSBs and SSBs) from multiple sources, ensuring accurate classification by referencing relevant studies. From these annotated proteins, we derived sequence embeddings using pre-trained language models, including ESM2 [32], ESM1v [33], and ProtTrans [34], which capture intricate sequence features. Leveraging these embeddings, we constructed a deep learning model with a multi-channel input architecture. Through ablation experiments, we optimized the model’s configuration to maximize its predictive performance. Our model demonstrates superior performance in predicting ΔΔG for mutations in both DSBs and SSBs, outperforming existing state-of-the-art methods.

## 2. Materials and Methods

### 2.1 Benchmark Datasets Construction

To investigate the effects of missense mutations on protein-DNA interactions, we constructed a dataset capturing ΔΔG from 324 mutations across 73 complexes. This collection draws on foundational research, notablyPEMPNI [20], mCSM-NA [19] and SAMPDI [11], which sourced data from dbAMEPNI [17] and ProNIT [17]. Each entry specifies the mutation position, amino acid replacement, and associated ΔΔG value. For precision, we corroborated these mutation sites with sequence records from the Protein Data Bank [35] (as depicted in **Figure 1A-1B**). Following PEMPNI’s methodology, the dataset was divided into a training set, MPD276 (276 mutations from 53 complexes), and an independent test set, MPD48 (48 mutations from 20 complexes), with MPD48 excluding data utilized by the referenced algorithms. Subsequently, proteins in both sets were categorized into DSBs and SSBs using ground truth provided by iPNHOT [22, 28], detailed information is provided in **Table S1** and **Table S2**.

**Figure 1.**
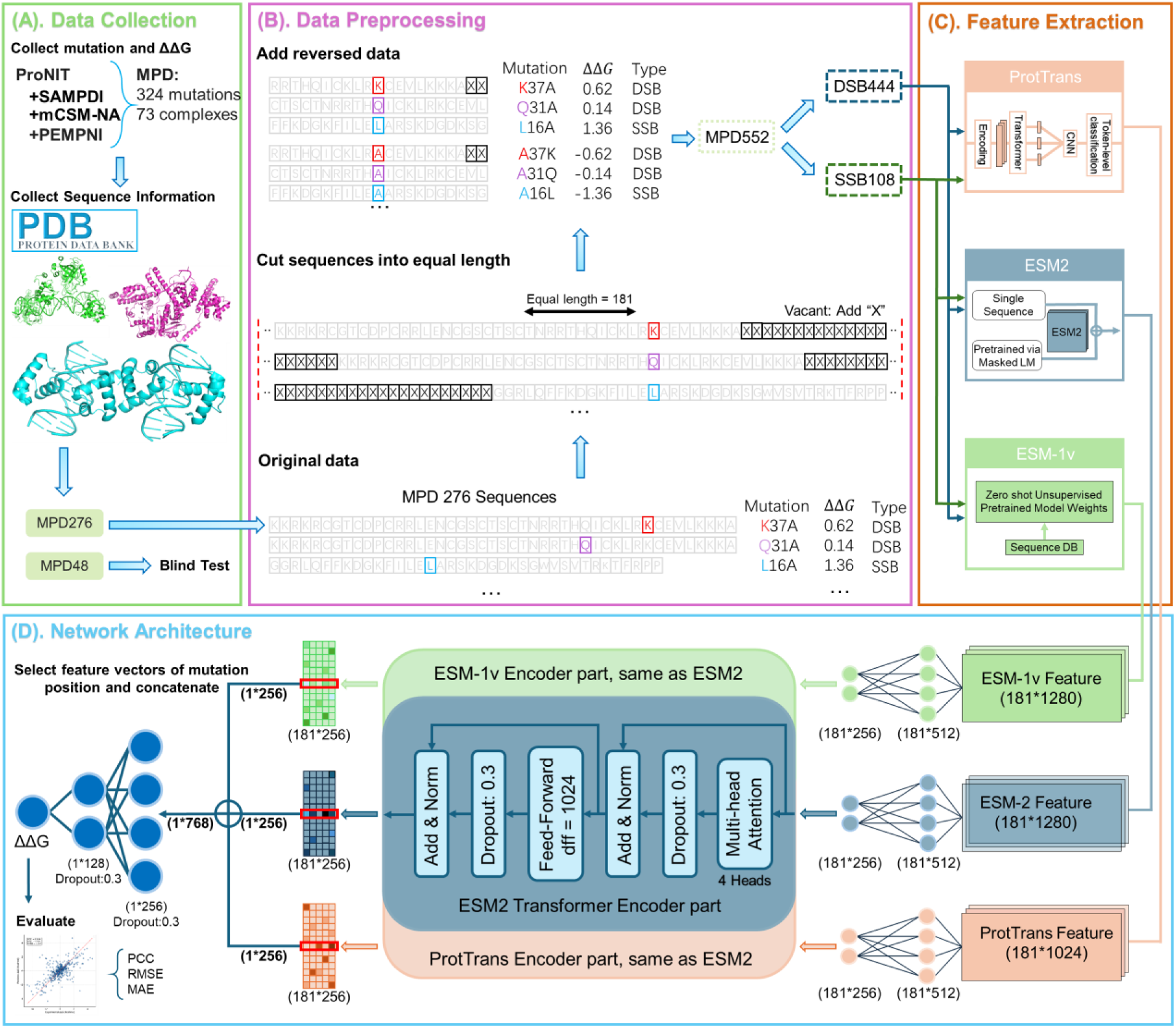
Overview of the EPEPDI workflow. **(A)** Mutation data collection; **(B)** Mutation data preprocessing; (C) Feature Extraction; **(D)** Architecture of EPEPDI.

As outlined in **Table 1**, MPD276 consists of DSB222 (222 mutations from 47 complexes) and SSB54 (54 mutations from 6 complexes), while MPD48 comprises IDSB35 (35 mutations from 15 complexes) and ISSB13 (13 mutations from 5 complexes). To ensure a more balanced dataset, reverse mutations were introduced, guided by the thermodynamic principle that the free energy change (ΔΔG) for a mutation from wild type to mutant equals, in magnitude but opposes in sign, that of the reverse mutation (i.e., ΔΔG_wildtype→mutant_=ΔΔG_mutant→wildtype_). This refinement is evident in the revised notations, such as MPD552 and DSB444, as documented in **Table 1**.

**Table 1.** Statistical summary of benchmark datasets.

| Dataset Name | Complexes | Mutations | Notation (including Reversed $\Delta\Delta G$) |
| --- | --- | --- | --- |
| MPD276 | 53 | 276 | MPD552 |
| DSB222 | 47 | 222 | DSB444 |
| SSB54 | 6 | 54 | SSB108 |
| MPD48 | 20 | 48 | MPD96 |
| IDSB35 | 15 | 35 | IDSB70 |
| ISSB13 | 5 | 13 | ISSB26 |

### 2.2 Feature Representation

Robust numerical feature extraction from protein sequences is vital for deep learning models designed to predict the effects of missense mutations [36]. This process enables models to uncover intricate patterns within protein structures. Advanced protein language models (PLMs), including ProtTrans [34], ESM-2 [32], and ESM-1v [33], leverage extensive training datasets to produce per-residue embeddings that encapsulate critical protein characteristics, particularly around mutation sites (as depicted in **Figure 1C**).

ProtTrans [34], developed using the BFD dataset with 393 billion amino acids [37, 38], leverages 1024-dimensional embeddings per residue. For this study, we standardized sequences to 181 residues, centered on the mutation site, producing a 181×1024 feature matrix per sample. These embeddings adeptly reflect protein structure and function constraints, proven effective in tasks like secondary structure prediction [39].

In contrast, ESM-2 [32], a Transformer model with 650 million parameters, leverages 65 million distinct UniRef sequences [31, 40] through masked language modeling [41, 42]. This approach enables it to discern complex sequence patterns, producing 1280-dimensional per-residue embeddings. Applied to our 181-residue sequences, it produces a 181×1280 matrix, providing deep insights into the mutation site’s local environment.

Similarly, ESM-1v [33], also featuring 650 million parameters, excels at zero-shot variant effect prediction. Trained on 98 million unannotated sequences [32, 43], it generates 1280-dimensional embeddings per residue, forming a 181×1280 matrix for each 181-residue sequence. This representation adeptly reflects evolutionary and contextual details [33].

To ensure uniformity, we fixed the sequence length at 181 residues across all models, a decision guided by entropy analysis (please refer to **Section 4.2** for more details) to optimize both predictive performance and computational efficiency. Sequences were aligned at the mutation site, with padding or truncation implemented as required. For the MPD552 dataset, we assembled feature tensors—such as 512×181×1280 for ESM-2—omitting batch dimensions for simplicity/.

### 2.3 Performance Evaluation Methods

In this study, the dataset was split into 90% for training (with 10% of the training data randomly assigned as a validation set) and 10% for testing. The model was trained over 150 epochs, during which the validation set tracked performance, improvements triggered model saving. To avoid overfitting, early stopping halted training if validation loss stalled for 40 epochs. For robust assessment, ten-fold cross-validation divided the DSB444 dataset into ten equal parts, iteratively using nine for training and one for testing, with averaged metrics ensuring reliable generalization estimates. Finally, independent datasets (MPD96 for the full model, IDSB70 for DSB-focused model, and ISSB26 for SSB-focused model), served as blind tests to evaluate performance across diverse protein types.

### 2.4 Evaluation Metrics

To assess the EPEPDI model’s predictive accuracy, we combined multiple metrics to capture distinct performance aspects. The Pearson Correlation Coefficient (PCC) to measure trend consistency between predicted and true values. However, PCC falls short in assessing absolute accuracy, as it may remain high despite consistent offsets in predictions. To overcome this, we incorporate the Root Mean Square Error (RMSE), which amplifies larger deviations, thus highlighting significant errors, and the Mean Absolute Error (MAE), which offers a straightforward, robust average error metric less swayed by outliers. Together, these metrics provide a thorough evaluation of the EPEPDI model’s performance and enable comparisons with established methods. Specifically, PCC tracks correlation, while RMSE (kcal/mol) and MAE (kcal/mol) gauge prediction precision against experimental data (Detailed metric descriptions are available in **Text S1**).

## 3. EPEPDI Network Architecture

In this study, we present EPEPDI, a novel computational model designed to predict alterations in ΔΔG caused by missense mutations in PDIs (**Figure 1D)**. EPEPDI utilizes three distinct feature matrices derived from PLMs: ESM-1v, ESM-2, and ProtTrans, each centered on the mutation site.

Initially, these features undergo dimensionality reduction through two fully connected layers, yielding compact 181×256 representations. Positional encoding is then applied to preserve sequence order. Subsequently, three dedicated transformer encoders, each comprising two layers with multi-head attention (4 heads) and feedforward networks, process these encoded features. This design enables the model to capture intricate contextual dependencies, while dropout, layer normalization, and residual connections enhance training stability and mitigate overfitting.

Next, feature vectors at the mutation site are extracted from each encoder’s output and concatenated into a 768-dimensional vector. This combined vector is fed into a neural network with two hidden layers (256 and 128 units) and an output layer that delivers the predicted ΔΔG. To optimize the model, we implemented the Pseudo-Huber loss function to assess the divergence between predicted *y*_pred_ and true values *y*_true_, defined as follows:

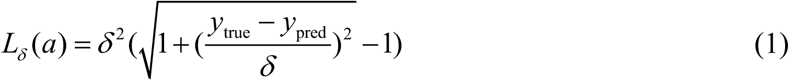

where *δ* is a hyperparameter, balancing precision for small errors with robustness against larger deviations. Multi-channel architecture in EPEPDI could integrate diverse protein representations, achieving high accuracy in predicting mutation effects on PDIs.

## 4. Results and Discussions

### 4.1 Comparison of DSBs and SSBs from Different Viewpoints

In this section, a systematic comparison of DSBs and SSBs was performed using the DSB222 and SSB54 datasets, as depicted in **Figure 2**. Analysis of ΔΔG distributions (**Figure 2A**) indicates that SSBs exhibit higher mean values and predominantly positive ΔΔG samples, whereas DSBs present a more even split between positive and negative values. Furthermore, bar charts of average ΔΔG across the 20 amino acids (**Figure 2B**) reveal notable differences at positions A, C, E, N, and P, with little overlap between DSBs and SSBs across mutation sites.

**Figure 2.**
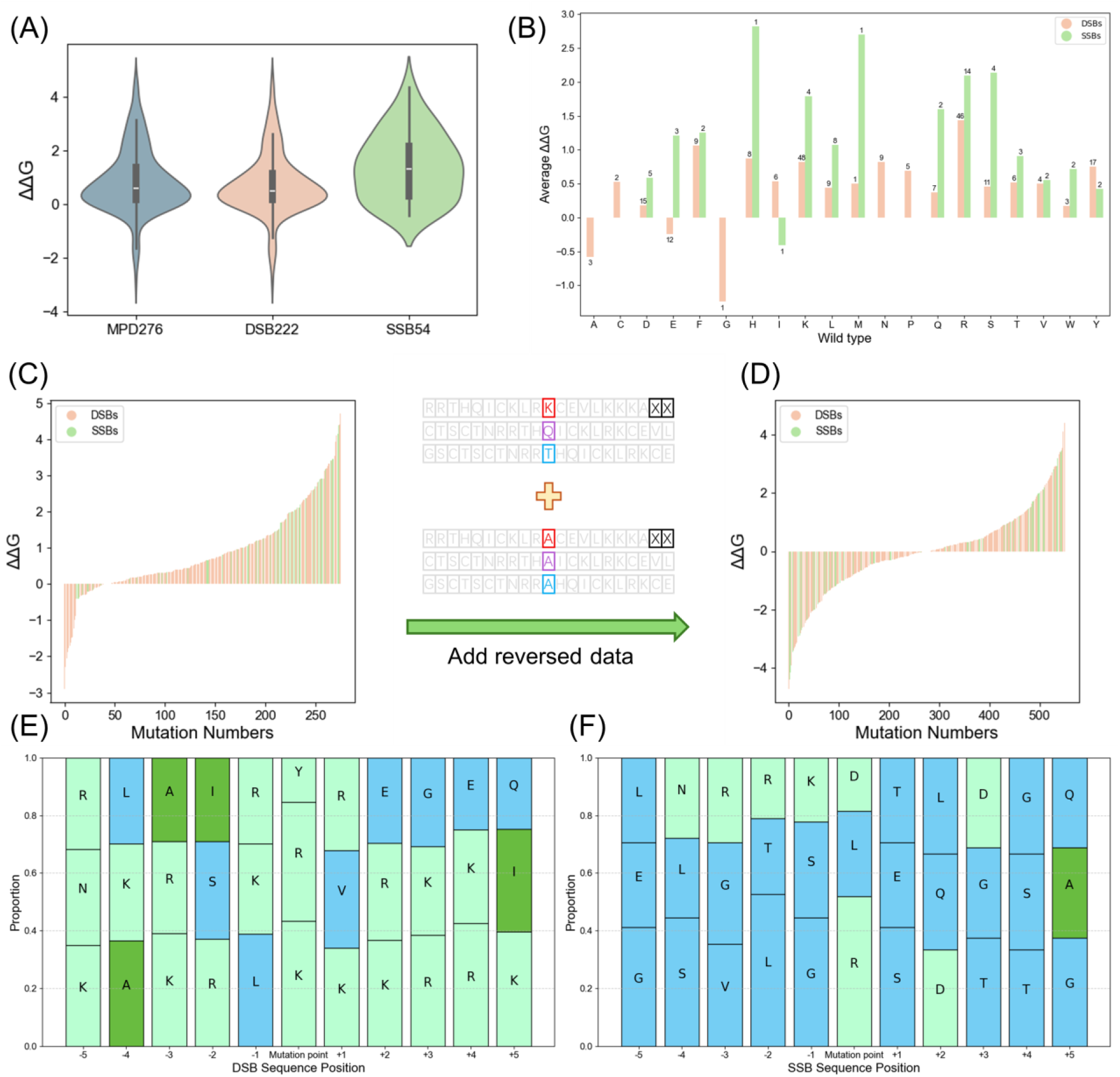
Comparative analysis of DSBs and SSBs sub-datasets. **(A)** ΔΔG distributions for MPD276, DSB222, and SSB54 datasets, revealing dataset-specific mutation effects; **(B)** Mean ΔΔG values across 20 amino acid mutation types in DSBs and SSBs, underscoring differential impacts; **(C)** Pre-augmentation ΔΔG distribution, skewed toward positive values, with a schematic depicting reverse augmentation (mutants revert to wild-type, ΔΔG negated); **(D)** Post-augmentation ΔΔG distribution, equilibrated with positive and negative values, optimizing deep learning applicability; **(E)** Top three amino acid frequencies within a ±5 residue window of DSBs mutation sites; **(F)** Top three amino acid frequencies within a ±5 residue window of SSBs mutation sites.

To address the skewed ΔΔG distribution (**Figure 2C**), a reverse augmentation approach was applied, reintroducing wild-type residues with negated ΔΔG values, yielding a balanced post-augmentation (**Figure 2D**) suitable for deep learning. Additionally, amino acid frequencies within a ten-residue window around mutation sites (**Figure 2E-2F**) show distinct profiles: DSBs are enriched with R and K, while SSBs favor G, T, and L. These contrasting sequence contexts imply significant variation in underlying data properties, potentially affecting feature extraction in machine learning models.

Consequently, given the pronounced differences in ΔΔG patterns and local sequence environments, training a single model on combined DSBs and SSBs data could impair predictive accuracy. Therefore, it is recommended that separate models be developed for each type of DNA binding protein to optimize performance and better capture their unique mutation effects.

### 4.2 Information Entropy Analysis for Different Input Sequence Length

When examining protein binding data, accounting for sequence length variation is essential. Typically, sequences are aligned around the missense mutation, trimmed to a uniform length (e.g., 41 AAs), and padded with ‘X’ [36, 44]. Selecting an optimal length is vital, as it significantly influences predictive model outcomes. Given that protein sequences resemble character strings, they align well with PLMs. To address this, we devised an algorithm that utilizes information entropy dynamics to determine the optimal sequence length.

#### Algorithm Calculate the entropy of data given linearly increasing sequence length values

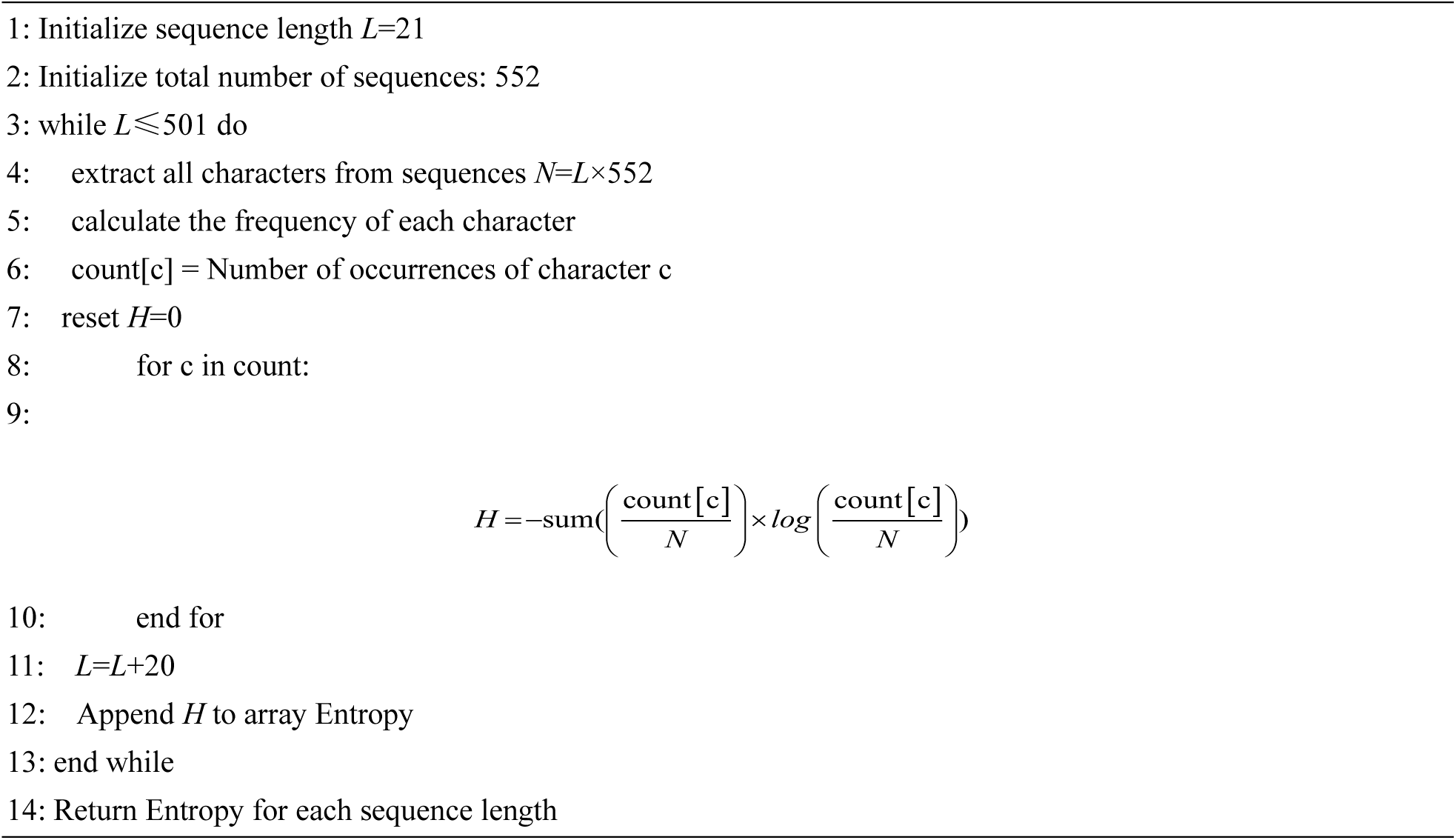

Our algorithm adjusts sequence length *L* from 21 to 501 in increments of 20, evaluating 552 sequences per *L*. It computes Shannon entropy using *H* formula in algorithm, in which, *count[c]* denotes the frequency of character *c*, and *N*=*L*×552 reflects total characters, measuring distribution evenness. Data analysis reveals distinct trends. As shown in **Figure 3A**, the rate of incorporation of valid amino acids (excluding ‘X’) declines with increasing length, stabilizing near zero by 501 AAs. This suggests that beyond a certain threshold, additional length contributes minimal new information. Meanwhile, entropy rises from 21 to 41 AAs before steadily decreasing (**Figure 3B**). Model performance, evaluated via ten-fold cross-validation and averaged PCC scores (**Figure 3C**), generally improves with decreasing entropy. Notably, an exception occurs at 41 AAs, where performance dips despite higher entropy, corresponding to a positive entropy change rate (**Figure 3D**).

**Figure 3.**
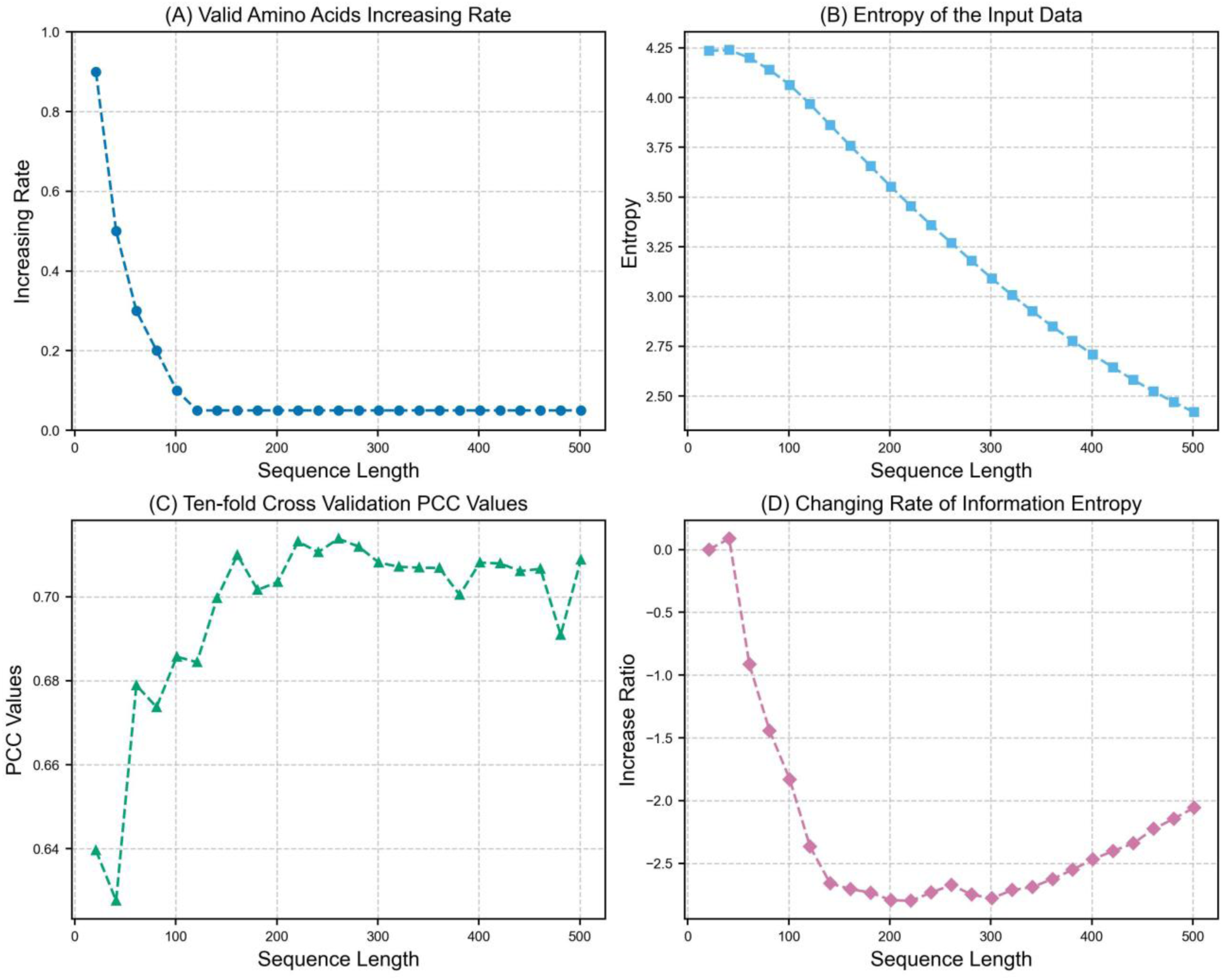
Analysis of sequence length effects on information metrics with sequence lengths from 21 to 501. **(A)** Valid amino acids increasing rate; **(B)** Entropy; **(C)** Ten-fold cross-validation PCC; **(D)** Changing rate of information entropy.

These results yield two key observations. First, predictive accuracy relies on adequate valid information; once this plateaus, performance gains cease. Second, high entropy may introduce noise, obscuring patterns, whereas excessively low entropy risks underfitting due to insufficient complexity. Consequently, we identified 181 AAs as the optimal length, where both the rate of valid amino acid increase and entropy change approach zero, striking a balance between accuracy and efficiency. This approach significantly refines our ability to predict mutation effects on DNA-binding proteins and establishes a versatile framework for broader protein dataset analyses.

### 4.3 EPEPDI Performances on Benchmark and Separated Datasets

To rigorously assess the EPEPDI model’s capability in predicting missense mutation effects on DNA-binding proteins, we conducted ten-fold cross-validation on three distinct datasets (including MPD552, DSB444, and SSB108) using uniform model parameters. The outcomes, detailed in **Table 2**, reveal varied performance across these sets. Specifically, on MPD552, EPEPDI attained an average PCC of 0.755, an MAE of 0.674, and an RMSE of 0.964 over ten folds. In contrast, training solely on DSB444 resulted in a diminished PCC of 0.681, a marginally elevated MAE of 0.694, and an RMSE of 1.031. Conversely, SSB108 training yielded a notable PCC increase to 0.863, accompanied by rises in MAE and RMSE to 0.764 and 0.9958, respectively. This indicates a PCC improvement of 0.108 for SSB108 and a decline of 0.074 for DSB444, underscoring the influence of dataset-specific traits. Since parameter optimization was not pursued, performance remained suboptimal, suggesting that tailored adjustments could enhance outcomes.

**Table 2.** Prediction performance of 10-fold cross-validation on benchmark datasets.

| Datasets | PCC (kcal/mol) | MAE (kcal/mol) | RMSE (kcal/mol) |
| --- | --- | --- | --- |
| MPD552 | 0.755±0.135 | 0.674±0.149 | 0.964±0.324 |
| DSB444 | 0.681±0.139 | 0.694±0.161 | 1.031±0.270 |
| SSB108 | 0.863±0.092 | 0.764±0.319 | 0.996±0.405 |

Further insights emerge from **Figure 4**, it demonstrates that MPD552 exhibits balanced performance across all metrics. Although DSB444’s PCC is lower, its MSE and RMSE display greater stability. For SSB108, despite a higher average PCC, performance fluctuates significantly across folds, likely due to its smaller size, where suboptimal data in any fold markedly affects results. These observations affirm EPEPDI’s promise for DNA-protein interaction prediction yet highlight the necessity for refinement when addressing imbalanced or limited datasets.

**Figure 4.**
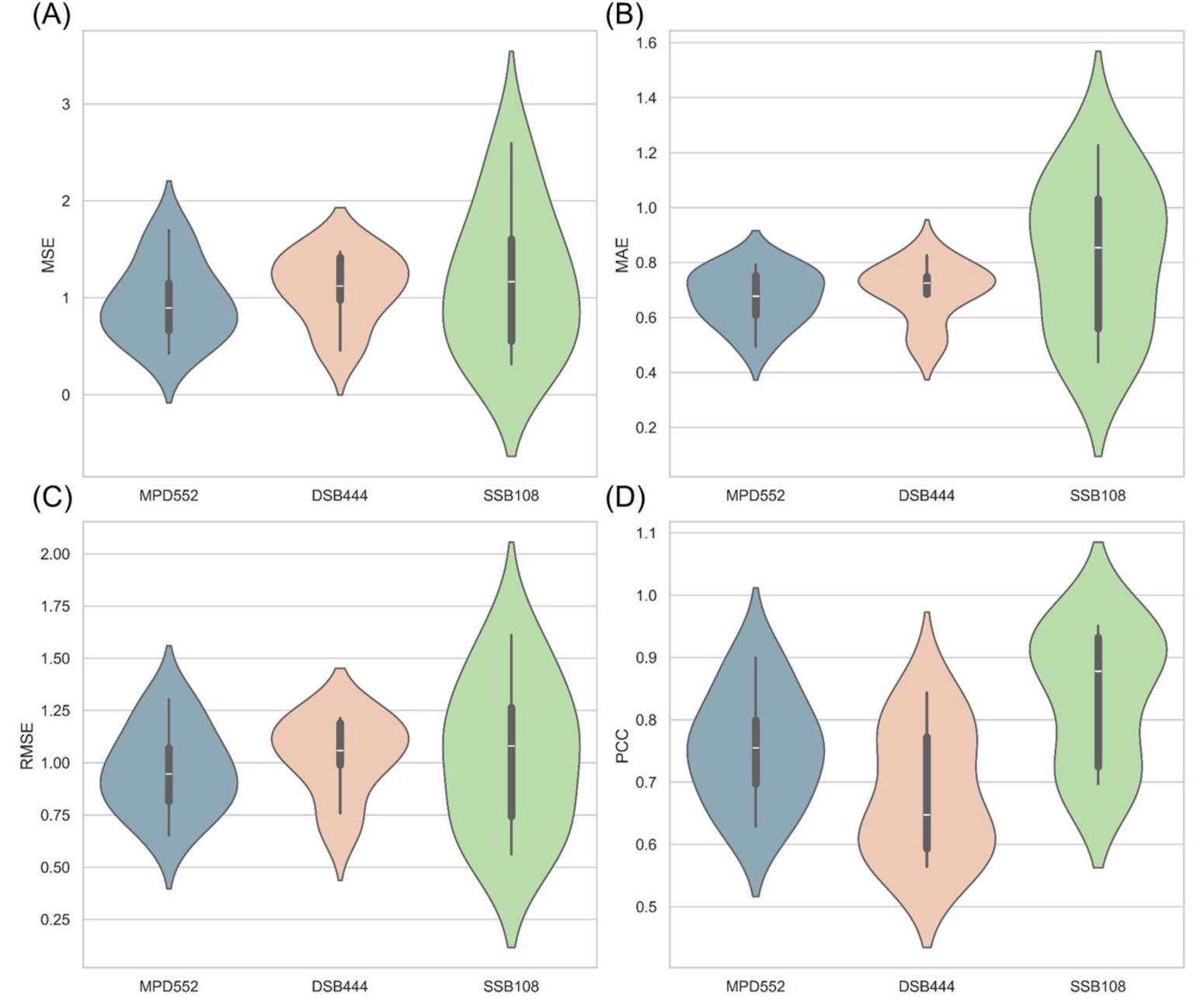
Performance evaluation of the EPEPDI model across datasets (including MPD552, DSB444, and SSB108) using ten-fold cross-validation. **(A)** MSE, **(B)** MAE, **(C)** RMSE, and **(D)** PCC.

### 4.4 Ablation Study

In our ablation study, we assessed the individual and combined contributions of three feature sets (i.e., ESM1v, ESM2, and ProtTrans) to the predictive performance of our model for missense mutations affecting DNA binding proteins. By training the model with different combinations of these features, we evaluated their impact using key metrics: PCC, MAE, and RMSE, with results averaged over ten-fold cross-validation (**Table 3**).

**Table 3.** Prediction performances of feature ablation studies on MPD552 dataset.

| Different Feature Set | PCC (kcal/mol) | MAE (kcal/mol) | RMSE (kcal/mol) |
| --- | --- | --- | --- |
| ESM1v | 0.731 | 0.667 | 1.027 |
| ESM2 | 0.687 | 0.736 | 1.089 |
| ProtTrans | 0.708 | 0.751 | 1.079 |
| ESM1v + ESM2 | 0.711 | 0.725 | 1.054 |
| ESM1v + ProtTrans | 0.713 | 0.712 | 1.052 |
| ESM2 + ProtTrans | 0.719 | 0.723 | 1.041 |
| ESM1v + ESM2 + ProtTrans | 0.755 | 0.674 | 0.964 |

The results reveal that the integration of all three feature sets yields the highest PCC (0.755) and the lowest RMSE (0.964), indicating a strong correlation between predicted and actual values and effective minimization of prediction errors. Notably, while the MAE for this comprehensive combination is marginally higher (0.674) compared to that of ESM1v alone (0.667), this slight increase is outweighed by the substantial enhancements in PCC and RMSE, thereby affirming the superiority of the full feature set.

Examining pairwise combinations further elucidates the feature interactions. For instance, the combination of ESM1v and ESM2 results in a PCC of 0.711, which is lower than that achieved by ESM1v individually (0.731). Parring ESM2 with ProtTrans produces a PCC of 0.719, exceeding the individual performances of both ESM2 (0.687) and ProtTrans (0.708), suggesting a synergistic effect between these two feature sets. Ultimately, the incorporation of all three features leads to the most pronounced performance improvement, underscoring the critical role of integrating diverse feature sets to achieve optimal predictive performance in modeling the effects of missense mutations on DNA binding proteins.

### 4.5 Performance Analysis of EPEPDI in Cross-Validation and Independent Testing

To thoroughly assess the predictive capabilities of EPEPDI, we systematically compared its performance against established methods using cross-validation and independent testing. This analysis draws on data from **Table 4**, **Figure 6**, and **Figure 5**. These sources provide a detailed basis for evaluating EPEPDI’s effectiveness and its distinctions from existing approaches.

**Figure 5.**
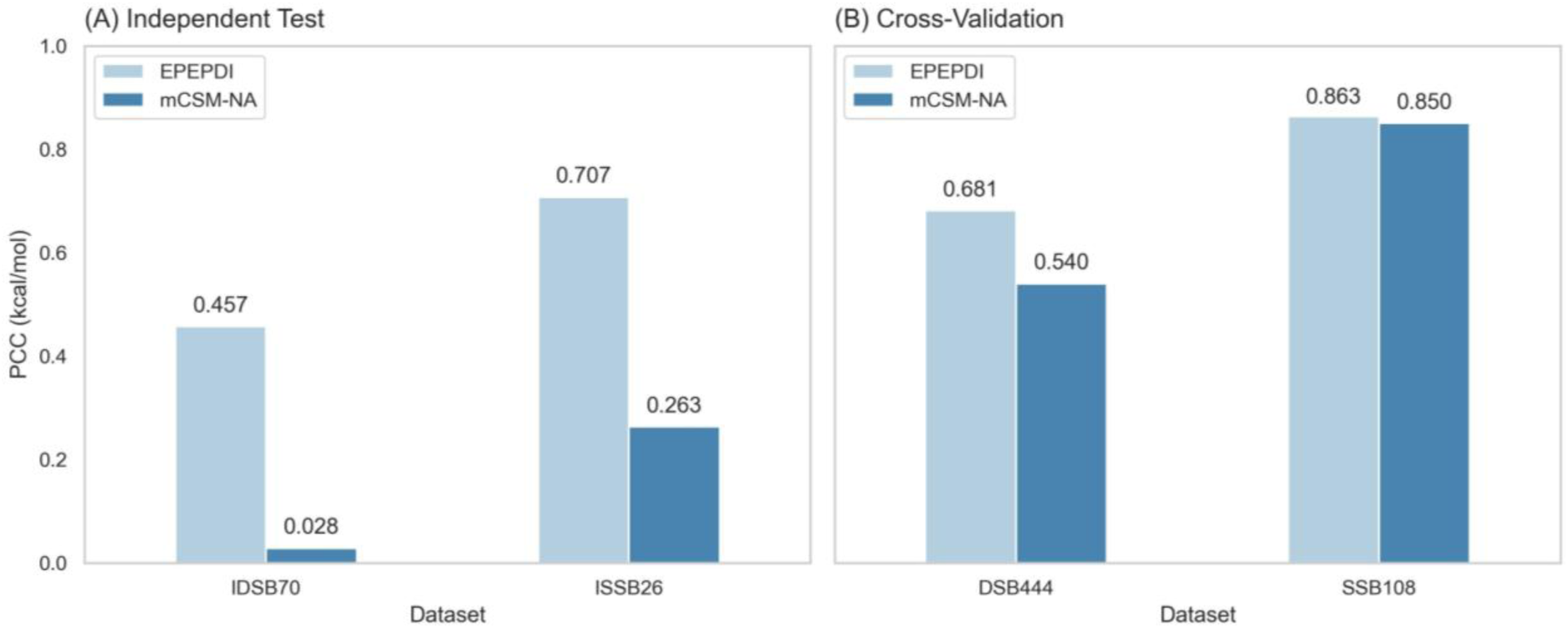
Comparison EPEPDI with mCSM-NA on DSB444, SSB108, DSB444, and SSB108 datasets.

**Figure 6.**
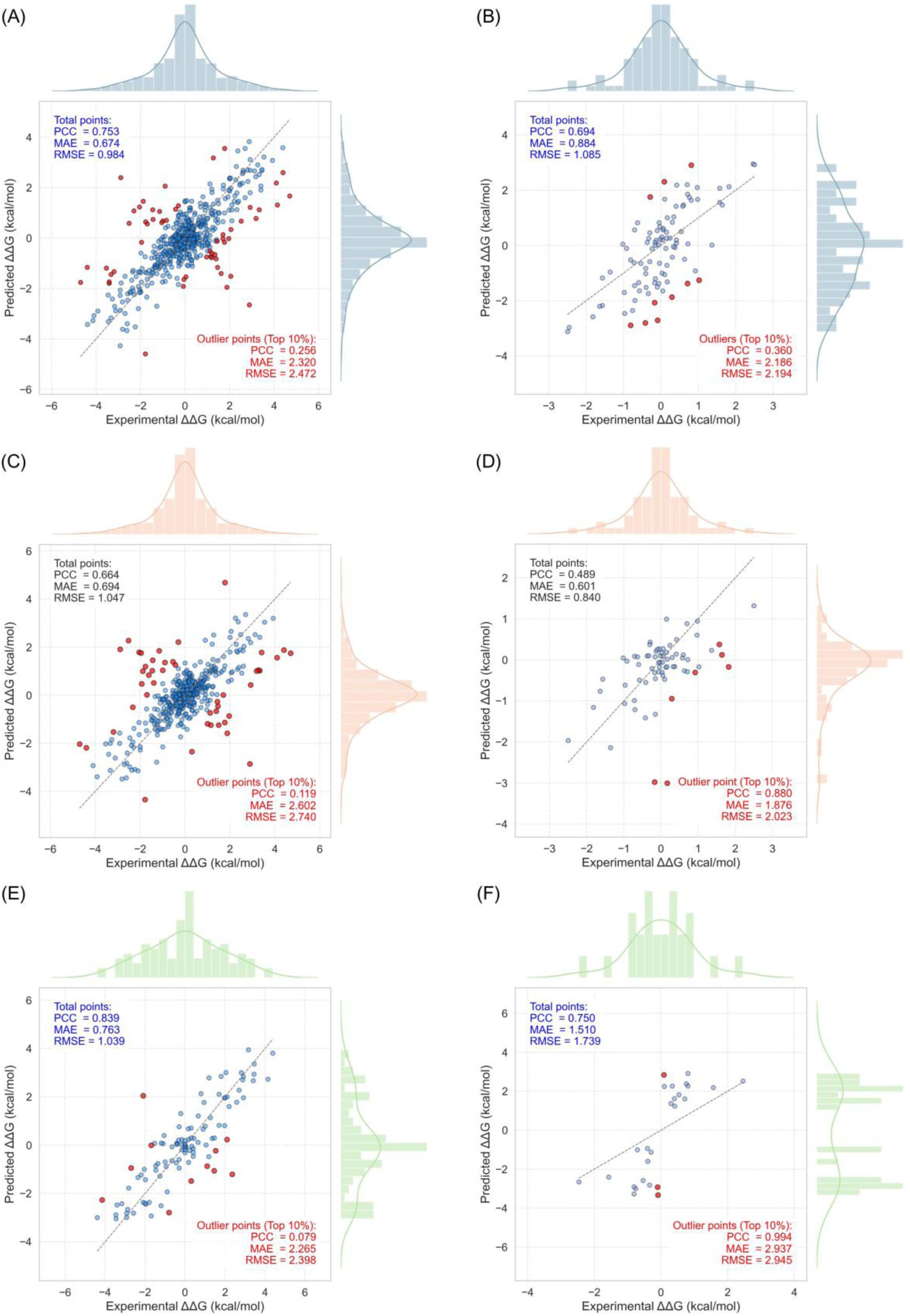
Scatter plot of different datasets. **(A)** MPD552 cross-validation, **(B)** MPD96 independent test, **(C)** DSB444 cross-validation, **(D)** IDSB70 independent test, **(E)** SSB108 cross-validation, and **(F)** ISSB26 independent test.

**Table 4.** Performance comparison with existing methods on MPD552 and MPD96 datasets Independent Test Set (MPD96)

| Validation Method and Dataset | Methods | PCC (kcal/mol) | RMSE (kcal/mol) |
| --- | --- | --- | --- |
| Cross-validation<br>(MPD552) | EPEPDI | 0.755 | 0.964 |
|  | PEMPNI | 0.575 | 0.978 |
| Independent Test Set<br>(MPD96) | EPEPDI | 0.632 | 0.942 |
|  | PEMPNI | 0.550 | 0.670 |
|  | SAMPDI | 0.424 | 0.705 |
|  | PremPDI | 0.509 | 0.887 |
|  | mCSM-NA | 0.016 | 1.259 |

#### 4.5.1 Performance Comparison on MPD552 and MPD96

As reported in **Table 4**, EPEPDI achieved an average PCC of 0.755 and an average RMSE of 0.964 during ten-fold cross-validation on the MPD552 dataset, surpassing PEMPNI (PCC = 0.575, RMSE = 0.978). This suggests greater predictive precision and consistency for EPEPDI on training data. Supporting this, **Figure 6A** presents a scatter plot for MPD552 cross-validation, showing a PCC of 0.753, a MAE of 0.674, and an RMSE of 0.883—values closely aligned with **Table 4**. For the scatter plots of models trained using ten-fold cross-validation, metrics are calculated by combining the results from each fold. For the scatter plots of models evaluated with the independent test, the best-performing model is selected for plotting. As a result, the values in the scatter plots may slightly differ from the average metrics presented in **Table 4**. The corresponding histogram indicates that both predicted and experimental values follow a near-normal distribution, largely within −2 to 2 kcal/mol, reinforcing the model’s reliability.

In independent testing on the MPD96 dataset, **Table 4** reveals EPEPDI’s average PCC of 0.632 and RMSE of 0.942, outperforming PEMPNI (PCC = 0.550, RMSE = 0.670), SAMPDI (PCC = 0.424, RMSE = 0.705), PremPDI (PCC = 0.509, RMSE = 0.887), and mCSM-NA (PCC = 0.016, RMSE = 1.259). EPEPDI’s notably higher PCC underscores its robust generalization to unseen data. However, PEMPNI’s lower RMSE (0.670) indicates a strength in limiting large prediction errors. **Figure 6B** further illustrates the superior performance of the best model among the ten-fold models on MPD96 independent test, reporting a PCC of 0.694, MAE of 0.884, and RMSE of 1.085—slightly differing from **Table 3**’s RMSE (0.942), possibly due to variations in data subsets or calculation methods, yet still affirming EPEPDI’s overall edge.

On the independent MPD96 test set, EPEPDI maintained a strong average PCC of 0.632, consistently exceeding other methods. Interestingly, its RMSE here is marginally lower than in cross-validation, reflecting stable performance. Since RMSE is sensitive to outliers, this metric highlights the model’s capacity to manage extreme values. For instance, **Figure 2**’s distribution plot of the MPD276 benchmark dataset reveals outliers like variant 2A0I_S3A (ΔΔG = 4.1 vs. series average 2.0) and 1AAY_R22A (ΔΔG = 3.54 vs. series average 1.19), far exceeding the dataset mean of 0.874. These anomalies emphasize EPEPDI’s generalization strength while explaining RMSE variations.

#### 4.5.2 Enhanced Performance via Differential DNA-Binder Training

Building upon core biophysical distinctions between dsDNA and ssDNA-binding proteins, we engineered a specialized training protocol for EPEPDI using DSB444 and SSB108 datasets. This structure-aware approach specifically accounts for differential binding mechanics. When benchmarked against mCSM-NA using independent test sets IDSB70 and ISSB26, **Figure 5** reveals striking PCC advantages. Complementing this, **Figure 6** delivers unprecedented diagnostic insights through integrated scatter plots that map prediction anomalies.

Notably, cross-validation exposes EPEPDI’s decisive edge: For DSB444, PCC jumps to 0.881 versus mCSM-NA’s 0.540 - a 63.3% relative gain. SSB108 shows similar dominance at 0.863 against 0.850. However, error decomposition uncovers critical nuances: While SSB108 maintains respectable aggregate accuracy (PCC=0.639, RMSE=1.039 kcal/mol), its extreme-value predictions collapse (top 10% outliers: PCC=0.079). This breakdown signals thermodynamic extrapolation failures at stability boundaries. DSB444 mirrors this vulnerability (outlier PCC=0.119) despite stronger baseline performance (PCC=0.664), highlighting persistent entropy modeling challenges in dsDNA recognition.

Independent testing yields fascinating contradictions: IDSB70’s outliers paradoxically approach perfection (PCC=0.896) while overall performance remains modest (PCC=0.489). This divergence suggests systematic residue-interface prediction errors rather than random noise. Conversely, ISSB26 achieves extraordinary outlier stability (PCC=0.994) alongside robust overall metrics (PCC=0.750, RMSE=1.739 kcal/mol). Collectively, these patterns demonstrate EPEPDI’s superior adaptability to SSB landscapes despite constrained training complexity. For DSBs however, performance plateaus reflect the high-dimensional conformational search space inherent to dsDNA recognition - particularly sequence-specific groove penetration and dynamic hydrogen-bonding arrays.

#### 4.5.3 Data Distribution and Model Validation

**Figure 6** visually validates EPEPDI’s performance through scatter plots and histograms. The scatter plots show predicted values aligning closely with experimental ones along the diagonal across datasets, indicating strong consistency. Histograms reveal a near-normal distribution for both predicted and experimental values, centered between −2 and 2 kcal/mol. Higher MAE (1.519) and RMSE (1.739) for ISSB26 likely reflect data variability. Post-reverse augmentation, MPD552’s value distribution balanced around a near-zero mean, amplifying the influence of outliers on RMSE and making it a sensitive error indicator. This reinforces EPEPDI’s robustness, especially on SSB data (PCC = 0.839 for SSB108, 0.750 for ISSB26).

EPEPDI excels over existing methods in PCC on MPD552 and MPD976 (**Table 3**). With separate DSB and SSB training, it outperforms mCSM-NA (**Figure 5**), as substantiated by scatter plots and histograms (**Figure 6**). Its prowess is most evident on SSB data (cross-validation PCC = 0.863, independent PCC = 0.707), though less pronounced on DSB data (cross-validation PCC = 0.681, independent PCC = 0.457), suggesting room for refinement targeting DSB-SSB structural disparities.

Overall, EPEPDI delivers exceptional accuracy and adaptability in predicting mutation-driven binding free energy shifts, outperforming peers across datasets and validation methods, thus providing a strong platform for future advancements.

#### 4.5.4 Performance Comparison with Different Machine Learning Predictors

Due to the limited size of the dataset used in this study, the original MPD276 dataset contains only 276 mutation entries. Even after applying reverse augmentation to form MPD552, the dataset comprises just 552 samples. Small-scale datasets often show better adaptability with machine learning algorithms. Therefore, we explored the fusion of three types of sequence-based features— ESM2, ESM1v, and ProtTrans via concatenation. The fused features were used as input to predict ΔΔG values through ten-fold cross-validation across various machine learning regressors, including Support Vector Regression [45], Random Forest [46], Gradient Boosting [47], AdaBoost [48], Gaussian Process Regression [49], and XGBoost [50].

As shown in **Table 5**, our proposed model, EPEPDI, consistently outperformed all machine learning-based regressors. Among the traditional methods, Support Vector Regression achieved the highest PCC of 0.451. In contrast, XGBoost performed the worst, with a PCC of 0.081, an RMSE of 1.697, and an MAE of 1.164, representing the lowest performance across all metrics. In comparison, EPEPDI attained a PCC of 0.755, an RMSE below 1, and an MAE of only 0.674. These results clearly demonstrate the superior performance of EPEPDI and underscore the necessity of deep learning approaches in addressing high-dimensional and complex feature representations. For detailed parameters of predictors shown in **Table 5**, please refer to **Table S3**.

**Table 5.** Performance comparison with different predictors on MPD552 using ten-fold cross-validation.

| Predictor | PCC (kcal/mol) | RMSE (kcal/mol) | MAE (kcal/mol) |
| --- | --- | --- | --- |
| Support Vector Regression | 0.451 | 1.551 | 1.138 |
| Random Forest | 0.314 | 1.615 | 1.127 |
| Gradient Boost | 0.264 | 1.560 | 1.096 |
| AdaBoost | 0.242 | 1.512 | 1.081 |
| Gaussian regression | 0.246 | 1.424 | 1.035 |
| XGBoost | 0.081 | 1.697 | 1.164 |
| **EPEPDI** | **0.755** | **0.964** | **0.674** |

#### 4.5.5 Feature Visualization Analysis Explaining the Strong Performance of EPEPDI

**Figure 7** employs t-SNE [51] to visualize embeddings from ESM2, ESM1v, and ProtTrans. Each point in these maps is colored according to its ΔΔG value, enabling an assessment of how effectively each model differentiates stabilizing from destabilizing mutations.

**Figure 7.**
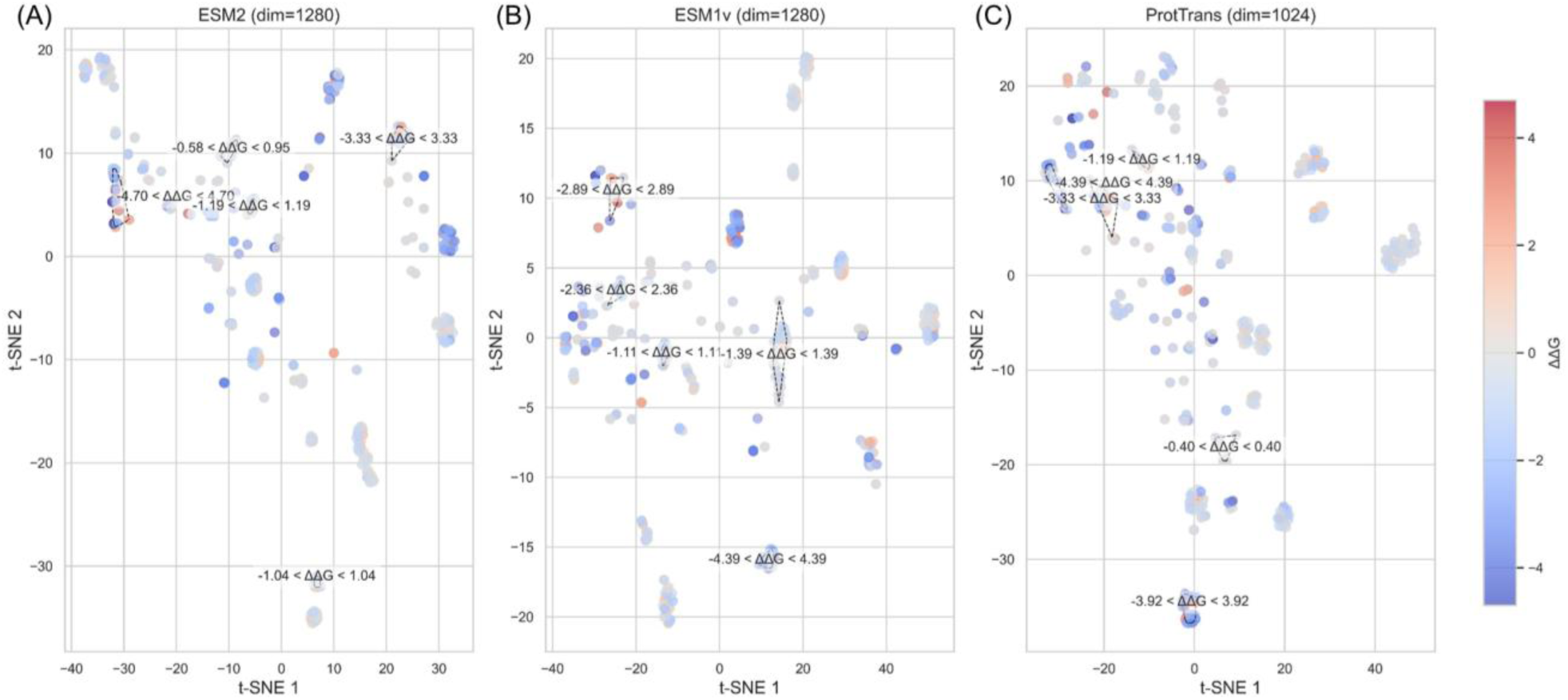
t-SNE map of protein large language model embeddings for discriminating ΔΔG values. **(A)**-**(C)** ESM2, ESM1v, and ProtTrans.

To analyze clustering behavior and discriminative strength, we applied DBSCAN [52] to the t-SNE coordinates. Here, the “eps” parameter was set as the median distance to the 15th nearest neighbor, and “min_samples” was fixed at 15 to detect dense clusters. For each subplot, we selected 3 to 5 representative clusters based on their mean ΔΔG values and evenly distributed indices, ensuring a broad representation across the ΔΔG range.

In **Figure 7A**, ESM2 embeddings yielded 15 clusters (eps = 2.19). Representative clusters spanned ΔΔG ranges such as −2.50 to −1.00 and 1.00 to 2.50, demonstrating clear separation between stabilizing and destabilizing mutations. In **Figure 7B**, ESM1v embeddings produced 12 clusters (eps = 2.33). However, some clusters showed overlapping ΔΔG ranges, suggesting less distinct separation. In **Figure 7C**, ProtTrans embeddings generated 13 clusters (eps = 3.35), capturing wider ΔΔG intervals, such as −3.00 to −1.50 and 0.50 to 3.00, which indicates robust discriminative ability.

The clustering analysis highlights model-specific ΔΔG patterns. For instance, ESM2 excels at resolving intermediate mutation effects (ΔΔG∈[−2.50, −1.00] and [1.00, 2.50]), while ProtTrans captures a broader range, including extreme values (e.g., [−3.00, −1.50] and [0.50, 3.00]), reflecting greater sensitivity to diverse mutational impacts. In contrast, ESM1v, with fewer clusters and some overlap, may provide specialized insights within narrower ΔΔG bands.

These distinct clustering profiles suggest that each model uncovers unique facets of the mutational landscape. Consequently, their embeddings could offer complementary perspectives for analyzing mutation effects. (For additional details on feature contributions, refer to **Table 3** and **Section 4.4**).

### 4.6 Case Study

This case study evaluates the predictive performance of the EPEPDI model, trained using ten-fold cross-validation on the MPD552 dataset, against the mCSM-NA [19] (through web server) across six mutation cases from three distinct proteins, as detailed in **Table 5**. The analysis examines mutations causing both substantial and minimal changes in ΔΔG, assessing performance for DSBs and SSBs DNA binding proteins. **Figure 8** complements this analysis by providing detailed visualizations of the local protein structures and mutation sites, highlighting their proximity to protein-DNA interfaces, which is critical for understanding their impact on binding affinity.

**Figure 8.**
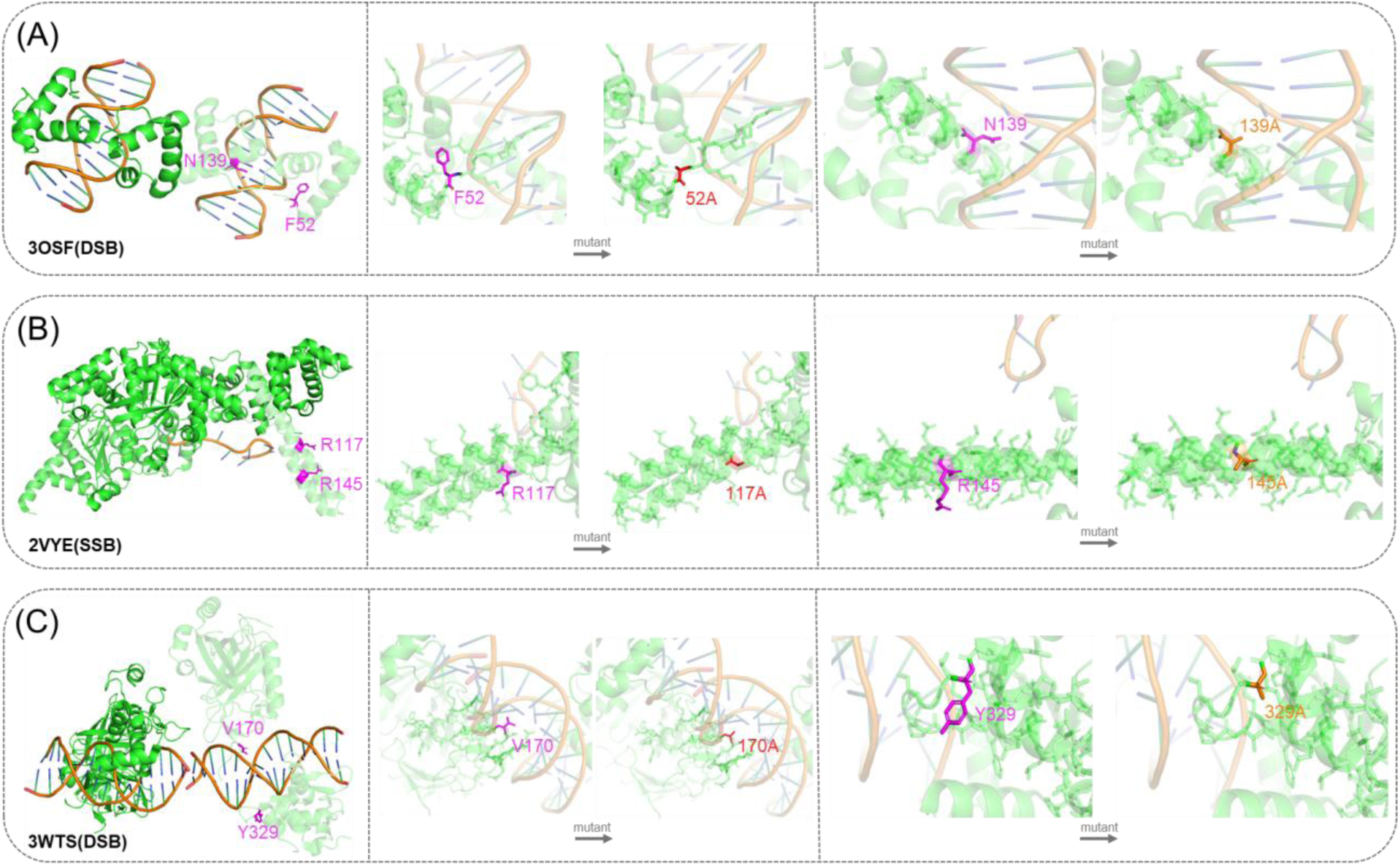
Protein local structure and mutation information. **(A)** PDB_ID: 3OSF (F52A, N139A). **(B)** PDB_ID: 2VYE (R117A, R145A). **(C)** PDB_ID: 3WTS (V170A, Y329A).

**Figure 8** displays the structures of three proteins (3OSF, 2VYE, and 3WTS) highlighting specific mutations. Each subplot displays a ribbon diagram of the protein’s overall structure and magnified views of both wild-type and mutant residues in stick form, emphasizing mutation sites proximate to protein-DNA interfaces. In **Figure 8A** (3OSF), mutations F52A and N139A are located at the interface, potentially disrupting essential DNA interactions. **Figure 8B** (2VYE) shows mutations R117A and R145A near the interface, which are likely to influence single-stranded DNA binding. **Figure 8C** (3WTS) presents mutations V170A and Y329A adjacent to a DNA molecule, suggesting their involvement in double-stranded DNA interactions. Specifically, in **Figure 8C** for the 3WTS protein, the V170A mutation is associated with a negligible change in binding affinity, as indicated by a ΔΔG value of 0.10 kcal/mol. Valine at position 170, being hydrophobic, is thought to contribute to the stability of the protein’s hydrophobic core or to engage in van der Waals interactions with nearby DNA regions. The mutation to alanine reduces the side-chain volume while maintaining hydrophobicity, resulting in only a minor effect on binding affinity.

As listed in **Table 6**, for mutations inducing significant changes in binding affinity (ΔΔG≥ 1.97 kcal/mol), EPEPDI exhibits superior predictive accuracy across both DSB and SSB proteins. For instance, in the DSB protein 3OSF, the N139A mutation, located at the protein-DNA interface (**Figure 8A**), has an experimental ΔΔG of 2.28 kcal/mol. EPEPDI predicts 2.31 kcal/mol, demonstrating excellent agreement, whereas mCSM-NA predicts −0.266 kcal/mol, incorrectly indicating a stabilizing effect. Similarly, the F52A mutation in 3OSF, also at the interface (**Figure 8A**), has an experimental ΔΔG of 2.81 kcal/mol, with EPEPDI predicting 2.703 kcal/mol, closely aligning with the true value. In contrast, mCSM-NA’s prediction of −1.391 kcal/mol is erroneous in both sign and magnitude

**Table 6.** Performance comparison with different predictors on MPD552 using ten-fold cross-validation.

| PDB_ID | Chain | Mutation | Type | $\Delta\Delta G$ (kcal/mol) | EPEPDI | mCSM-NA |
| --- | --- | --- | --- | --- | --- | --- |
| 3OSF | A | N139A | DSBs | 2.28 | 2.31 | -0.266 |
| 3OSF | A | F52A | DSBs | 2.81 | 2.703 | -1.391 |
| 2VYE | A | R145A | SSBs | 1.97 | 1.905 | -1.350 |
| 2VYE | A | R117A | SSBs | 2.00 | 1.862 | -1.351 |
| 3WTS | A | V170A | DSBs | 0.10 | 0.138 | -0.098 |
| 3WTS | C | Y329A | DSBs | 0.31 | 0.204 | -0.402 |

In the SSB-associated protein 2VYE, EPEPDI accurately predicts 1.905 kcal/mol for the R145A mutation (experimental ΔΔG = 1.97 kcal/mol) and 1.862 kcal/mol for the R117A mutation (experimental ΔΔG = 2.00 kcal/mol), both located near the DNA-binding interface (**Figure 8B**). Conversely, mCSM-NA yields negative predictions (−1.350 kcal/mol for R145A and −1.351 kcal/mol for R117A), failing to capture the destabilizing effect of these mutations. The structural proximity of these mutations to the DNA interface, as visualized in **Figure 8B**, likely contributes to their significant impact on binding affinity.

These results demonstrate that EPEPDI provides more precise and reliable predictions than mCSM-NA across both high and low ΔΔG mutations, achieving lower prediction errors for DSB and SSB datasets. The structural insights from **Figure 8** underscore the importance of mutation site locations near protein-DNA interfaces, which EPEPDI effectively captures. This enhanced accuracy positions EPEPDI as a robust tool for predicting the effects of missense mutations on DNA binding proteins.

## 5. Conclusions

PDIs play a pivotal role in fundamental biological processes such as DNA replication, repair, and gene regulation. Missense mutations in DNA-binding proteins can perturb these interactions by modifying binding affinities, which may lead to disease and hinder therapeutic progress. Accurately predicting the change in ΔΔG due to these mutations remains a significant challenge, primarily because current methods do not adequately address the structural variability of DNA-binding proteins and depend on hand-engineered features.

In response to these challenges, we introduce EPEPDI, an innovative two-stage deep learning framework that harnesses the power of transformer architecture and embeddings derived from ESM2, ESM1v, and ProtT5). A distinctive feature of EPEPDI is its ability to distinguish between DSBs and SSBs DNA-binding proteins, thereby facilitating accurate modeling of their unique binding characteristics. By concentrating on mutation sites, the framework effectively captures essential contextual information from both wild-type and mutant sequences.

Furthermore, we devised an information entropy-based algorithm to determine the optimal protein sequence length, establishing 181 residues as the point where information gain and entropy fluctuations reach equilibrium. This optimization not only improves predictive accuracy but also enhances computational efficiency and scalability. The framework’s first stage employs a transformer encoder with multi-head attention to amalgamate these features into a unified representation, which subsequently guides the ΔΔG prediction. EPEPDI demonstrates superior performance compared to existing state-of-the-art methods on a variety of benchmark and independent datasets, all while preserving computational efficiency.

## Author Contributions

Xuan Yu analysed the data, performed the experiments and wrote the original manuscript.

## Declaration of Competing Interest

The authors declare no conflict of interests.

## Notes

### Competing Interest Statement

The authors have declared no competing interest.

