## Supplementary Information for "EPEPDI: prediction of binding free energy changes from missense mutations in double and single-stranded DNA-binding proteins"

Xuan Yu[1]

[1]Department of Computer Science, City University of Hong Kong, Kowloon, Hong Kong, 999077, China.

**Supplementary Texts**

**Supplementary Text S1**

**Evaluation Metrics**

In regression analysis, we assess model accuracy using three key metrics: Pearson Correlation Coefficient (PCC) [1], Root Mean Square Error (RMSE) [2], and Mean Absolute Error (MAE) [2, 3].

PCC measures how well predicted and actual values align linearly, given by:

$$\rho = \frac{\mathrm{cov}(X,Y)}{\sigma_X \sigma_Y} \tag{1}$$

where $\mathrm{cov}(X,Y)$ is the covariance, and $\sigma_X$, $\sigma_Y$ are standard deviations. PCC values near 1 shows strong trend consistency, but it misses systematic shifts in predictions.

RMSE captures error magnitude, emphasizing large differences between true value and predicted value:

$$\mathrm{RMSE} = \sqrt{\frac{1}{n}\sum_{i=1}^{n}(y_i - \hat{y}_i)^2} \tag{2}$$

where $y_i$ denotes actual ΔΔG values and $\hat{y}_i$ as predicted ΔΔG values, RMSE (in kcal/mol) highlights outliers due to the squaring term.

MAE offers a simpler error average:

$$\mathrm{MAE} = \frac{1}{n}\sum_{i=1}^{n}|y_i - \hat{y}_i| \tag{3}$$

Also, in kcal/mol, MAE resists outlier influence, giving a steady accuracy view.

Together, PCC tracks correlation, while RMSE and MAE measure precision in binding affinity predictions for the EPEPDI model.

### Supplementary Tables

### Supplementary Table S1

Protein Type: double-stranded DNA binding proteins (DSBs) and single-stranded DNA binding proteins (SSBs).

**Table S1.** Ground truth of separating 47 types of DSBs and 6 types of SSBs in MPD276 benchmark dataset

| PDBID | Protein Names | Type |
|---|---|---|
| 1BP7 | MOBILE INTRON ENDONUCLEASE I-CREI | DSB |
| 1FOS | TWO HUMAN C-FOS:C-JUN | DSB |
| 1HCQ | THE ESTROGEN RECEPTOR DNA-BINDING DOMAIN | DSB |
| 2I05 | ESCHERICHIA COLI EPLICATIONTERMINATOR PROTEIN | DSB |
| 2MXF | The C-Terminal domain of MvaT | DSB |
| 2XRO | TtgV | DSB |
| 3ODC | HUMAN PARP-1 ZINC FINGER 2 | DSB |
| 3OSF | protozoan parasite Trichomonas vaginalis Myb2 in complex with MRE-2f-13 DNA | DSB |
| 3OSG | MYB-LIKE DNA-BINDING DOMAIN CONTAINING PROTEIN | DSB |
| 3QMG | CPG-BINDING PROTEIN | DSB |
| 3RN2 | INTERFERON-INDUCIBLE PROTEIN AIM2 | DSB |
| 3RN5 | AIM2 Inflammasome | DSB |
| 3RNU | Escherichia coli BL21(DE3) | DSB |
| 3SZQ | APRATAXIN-LIKE PROTEIN | DSB |
| 3UFD | Restriction-modification controller proteins | DSB |
| 3WTS | RUNT-RELATED TRANSCRIPTION FACTOR 1 | DSB |
| 4ATK | MITF:E-box complex | DSB |
| 4B5F | PUTATIVE EXODEOXYRIBONUCLEASE | DSB |
| 4BNC | HUMAN ETV1 | DSB |
| 4BXO | Fanconi anemia FANCM- FAAP24 complex | DSB |
| 4L0Z | Ets1 activation by Runx1 | DSB |
| 4L5R | INTERFERON-ACTIVABLE PROTEIN 202 | DSB |
| 4QJU | DNA-BINDING PROTEIN HU | DSB |
| 4QTJ | WHITE-OPAQUE REGULATOR 1 | DSB |
| 4R56 | CHROMATIN PROTEIN CREN7 | DSB |
| 4RDU | Dlx5 | DSB |
| 4XR0 | DNA REPLICATION TERMINUS SITE-BINDING PROTEIN | DSB |

| | | |
|---|---|---|
| 5D8C | MERR FAMILY REGULATOR PROTEIN | DSB |
| 5ED4 | RESPONSE REGULATOR | DSB |
| 5EXH | METHYLCYTOSINE DIOXYGENASE TET3 | DSB |
| 5KUB | DNA-7-METHYLGUANINE GLYCOSYLASE | DSB |
| 1AAY | ZIF268 ZINC FINGER | DSB |
| 1AIS | TATA-BINDING PROTEIN | DSB |
| 1AZ0 | ENDONUCLEASE ECORV | DSB |
| 1B3T | EBNA-1 NUCLEAR PROTEIN | DSB |
| 1B72 | HOMEOBOX PROTEIN HOX-B1 | DSB |
| 1CKQ | PRE-TRANSITION STATE ECO RI ENDONUCLEASE | DSB |
| 1CKT | HMG1 DOMAIN A BOUND TO A CISPLATIN-MODIFIED DNA DUPLEX | DSB |
| 1EWQ | DNA MISMATCH REPAIR PROTEIN MUTS | DSB |
| 1J5N | NONHISTONE CHROMOSOMAL PROTEIN 6A | DSB |
| 1MSE | C-MYB DNA-BINDING DOMAIN | DSB |
| 1PUF | HoxA9 and Pbx1 homeodomains bound to DNA | DSB |
| 1RUN | CATABOLITE GENE ACTIVATOR PROTEIN (CAP) | DSB |
| 1TN9 | TN916 INTEGRASE N-TERMINAL DOMAIN | DSB |
| 1TRO | TRP REPRESSOR OPERATOR COMPLEX | DSB |
| 3PVV | CHROMOSOMAL REPLICATION INITIATOR PROTEIN DNAA | DSB |
| 3VOK | Wild Type HrtR in the Apo Form with the Target DNA | DSB |
| 4WCG | ORF112 | SSB |
| 4ZSF | BSAWI ENDONUCLEASE | SSB |
| 5E24 | Su(H)-Hairless-DNA Repressor Complex | SSB |
| 1QZG | PROTECTION OF TELOMERES PROTEIN 1 | SSB |
| 2A0I | DNA HELICASE I | SSB |
| 2VYE | REPLICATIVE DNA HELICASE | SSB |

**Supplementary Table S2**

Protein Type: double-stranded DNA binding proteins (DSBs) and single-stranded DNA binding proteins (SSBs).

**Table S2.** Ground truth of separating 15 types of DSBs and 5 types of SSBs in MPD48 independent test set

| PDBID | Protein Names | Type |
|---|---|---|
| 4X5V | DNA POLYMERASE LAMBDA | DSB |
| 4XQ8 | DNA POLYMERASE LAMBDA | DSB |
| 4G4O | M77A mutation bound to oxoG-containing DNA | DSB |
| 4G4R | F114A mutation bound to oxoG-containing DNA | DSB |
| 3NGI | RB69 DNA Polymerase (Y567A) Ternary Complex with dTTP Opposite dG | DSB |
| 3OD8 | HUMAN PARP-1 ZINC FINGER 1 | DSB |
| 3SQ2 | RB69 DNA Polymerase Ternary Complex with dTTP Opposite 2AP | DSB |
| 4DU1 | RB69 DNA Polymerase Ternary Complex with dATP Opposite dT | DSB |
| 4HF1 | HTH-TYPE TRANSCRIPTIONAL REGULATOR ISCR | DSB |
| 4HN5 | HTH-TYPE TRANSCRIPTIONAL REGULATOR ISCR | DSB |
| 4NM6 | TET2-DNA complex | DSB |
| 5CO8 | NUCLEASE-LIKE PROTEIN | DSB |
| 5DFF | DNA-(APURINIC OR APYRIMIDINIC SITE) LYASE | DSB |
| 5DWB | TYPE-2 RESTRICTION ENZYME AGEI | DSB |
| 5U2R | DNA POLYMERASE BETA | DSB |
| 3WPD | horse TLR9 in complex with inhibitory DNA4084 | SSB |
| 4HQB | SINGLE-STRANDED DNA-BINDING PROTEIN DDRB | SSB |
| 5F55 | SINGLE-STRANDED-DNA-SPECIFIC EXONUCLEASE | SSB |
| 5IIN | pre-catalytic ternary extension complex of DNA polymerase lambda | SSB |
| 4K4G | DNA POLYMERASE LAMBDA IN COMPLEX WITH DNA AND L-DCTP | SSB |

### Supplementary Table S3

**Table S3.** Key hyperparameters of each predictor trained on MPD552 using ten-fold cross-validation.

| Predictor | Python Hyperparameter settings |
|---|---|
| Support Vector Regression | C=20, degree=1, gamma='auto' |
| Random Forest | max_depth = 10, min_samples_split = 5 |
| Gradient Boost | learning_rate = 0.01, max_depth = 8, n_estimators =100, subsample = 0.8 |
| AdaBoost | learning_rate = 0.1, n_estimators = 400 |
| Gaussian Regression | Constant kernel (1.0, (1e-4, 1e1)) * RBF kernel (1.0, (1e-4, 1e1)) |
| XGBoost | gamma = 0.1, learning_rate = 0.05, max_depth = 4, n_estimators = 300 |

### Supplementary Figure S1.

**Figure S1. Scatter plots of predicted vs. true labels across ten-fold cross-validation on DSDNA dataset.**

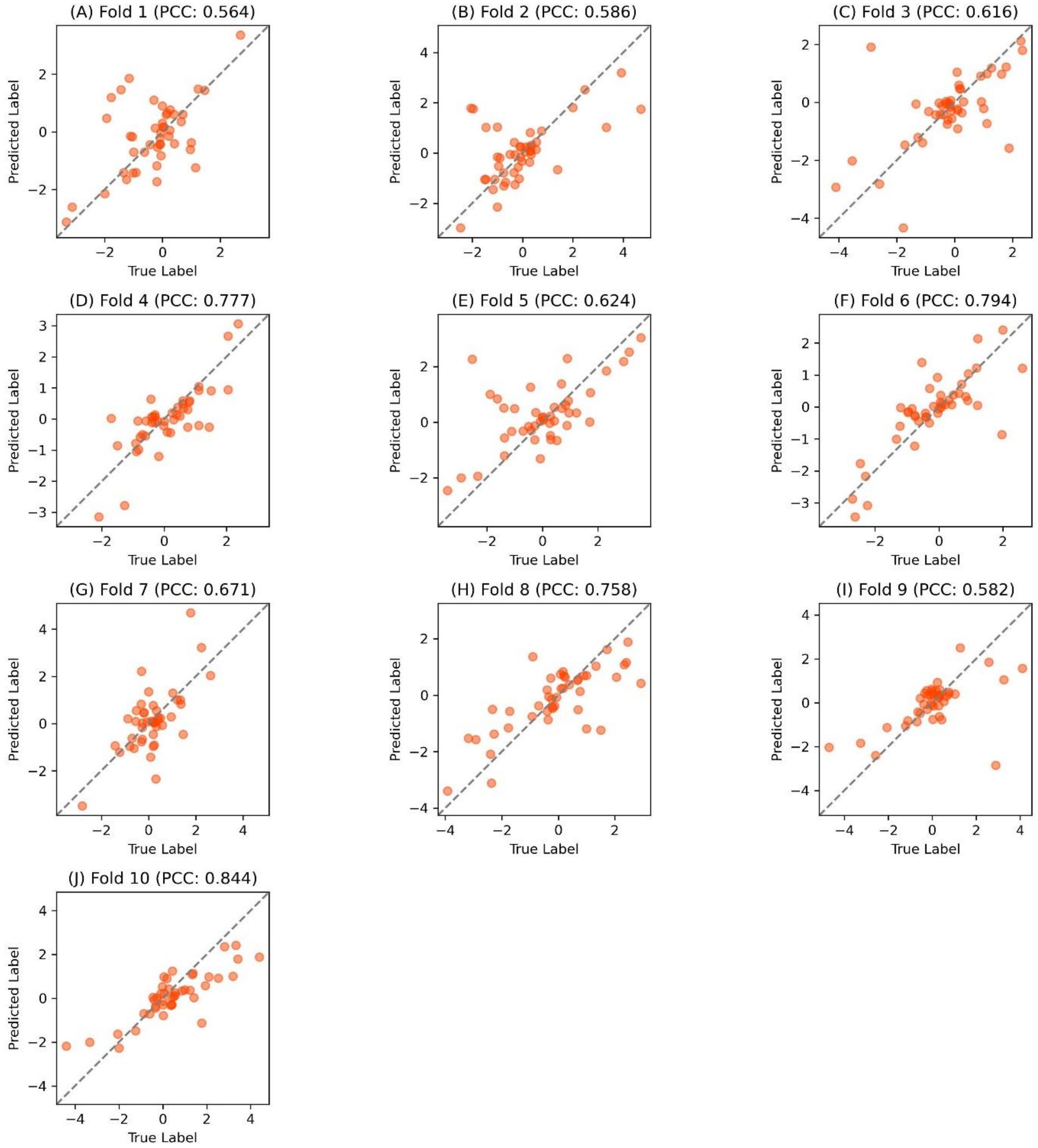


**Figure S1**. Scatter plots of predicted vs. true labels across ten-fold cross-validation on DSDNA dataset. (A) Fold 1 (PCC: 0.564), (B) Fold 2 (PCC: 0.586), (C) Fold 3 (PCC: 0.616), (D) Fold 4 (PCC: 0.777), (E) Fold 5 (PCC: 0.624), (F) Fold 6 (PCC: 0.794), (G) Fold 7 (PCC: 0.671), (H) Fold 8 (PCC: 0.758), (I) Fold 9 (PCC: 0.582), (J) Fold 10 (PCC: 0.844). Figure S1 illustrates the relationship between predicted and true labels across ten-fold cross-validation through scatter plots on DSDNA dataset, with each subplot representing a distinct fold. The x-axis denotes true labels, and the y-axis represents predicted labels, both ranging from approximately -4 to 4. Orange dots depict individual data points, with a dashed line (y = x) indicating perfect prediction. The PCC, ranging from 0.564 (Fold 1) to 0.844 (Fold 10), quantifies the linear correlation strength per fold. Fold 10 exhibits the highest PCC (0.844), with points tightly clustered along the diagonal, reflecting superior predictive accuracy. Conversely, Fold 1 (PCC: 0.564) shows greater dispersion, indicating weaker performance. The PCC variability (0.564–0.844) across folds underscores our model's sensitivity to data subset differences, emphasizing the critical role of cross-validation in evaluating robustness and generalization. Higher PCC values (e.g., Fold 6: 0.794, Fold 4: 0.777) suggest consistent predictive capability, while lower values (e.g., Fold 9: 0.582) highlight potential limitations in specific subsets, offering insights into model stability and areas for refinement.

### Supplementary Figure S2.

**Figure S2. Performance metrics across ten-fold cross-validation for the DSDNA dataset.**

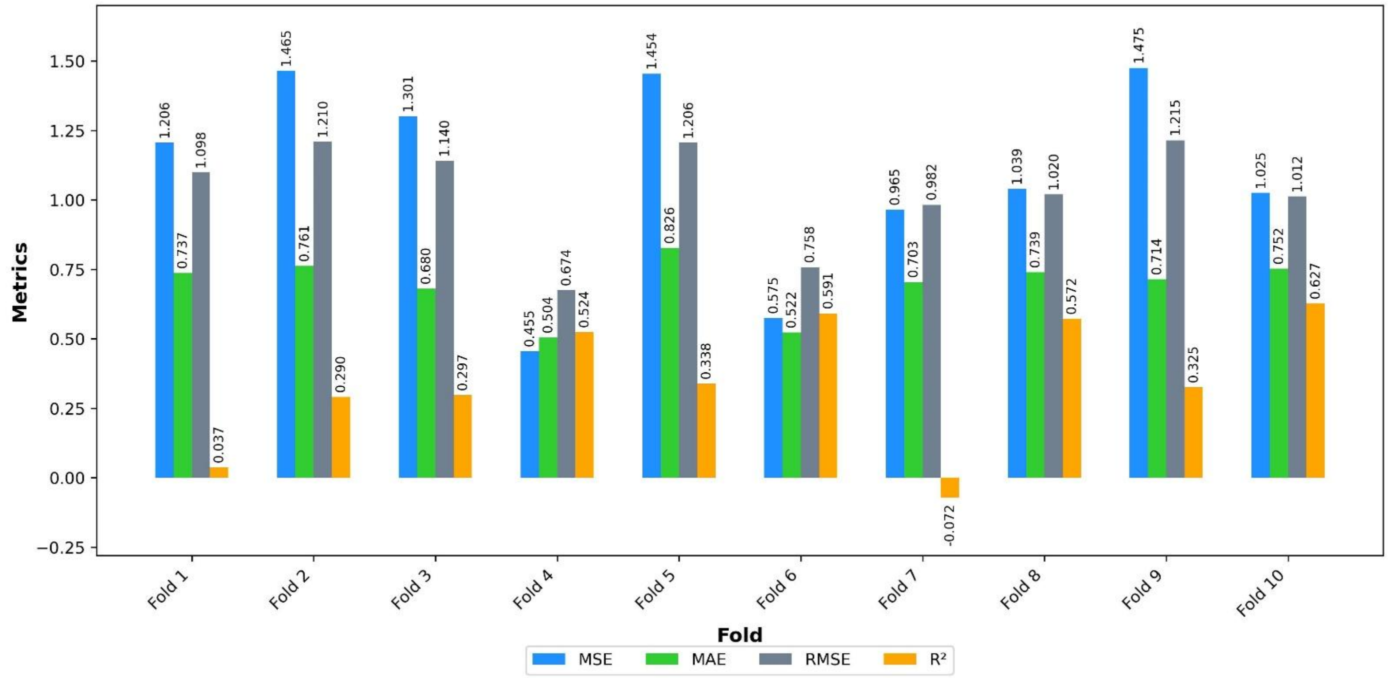


**Figure S2**. Performance metrics across ten-fold cross-validation for the DSDNA dataset.

Figure S2 provides a detailed assessment of the DSDNA model's performance across ten-fold cross-validation, revealing significant variability in predictive accuracy and explanatory power. Fold 4 demonstrates optimal performance with the lowest error metrics (MSE: 0.455, MAE: 0.504, RMSE: 0.674) and a robust $R^2$ (0.524), indicating strong predictive capability. Conversely, Fold 7 exhibits the weakest fit, with the highest errors (MSE: 0.965, MAE: 0.703, RMSE: 0.982) and a negative $R^2$ (-0.072), suggesting the model fails to explain variance in that subset. Fold 10 achieves the highest $R^2$ (0.627), reflecting superior explanatory power, consistent with its elevated PCC (0.844). The fluctuating $R^2$ and error metrics underscore the model's sensitivity to data partitioning, highlighting the necessity of cross-validation to evaluate robustness. These insights are pivotal for identifying performance disparities and guiding model optimization, emphasizing the interplay between error metrics and explanatory capacity in assessing reliability.

### Supplementary Figure S3.

#### Figure S3. Scatter plots of predicted vs. true labels across ten-fold cross-validation on the SSDNA dataset.

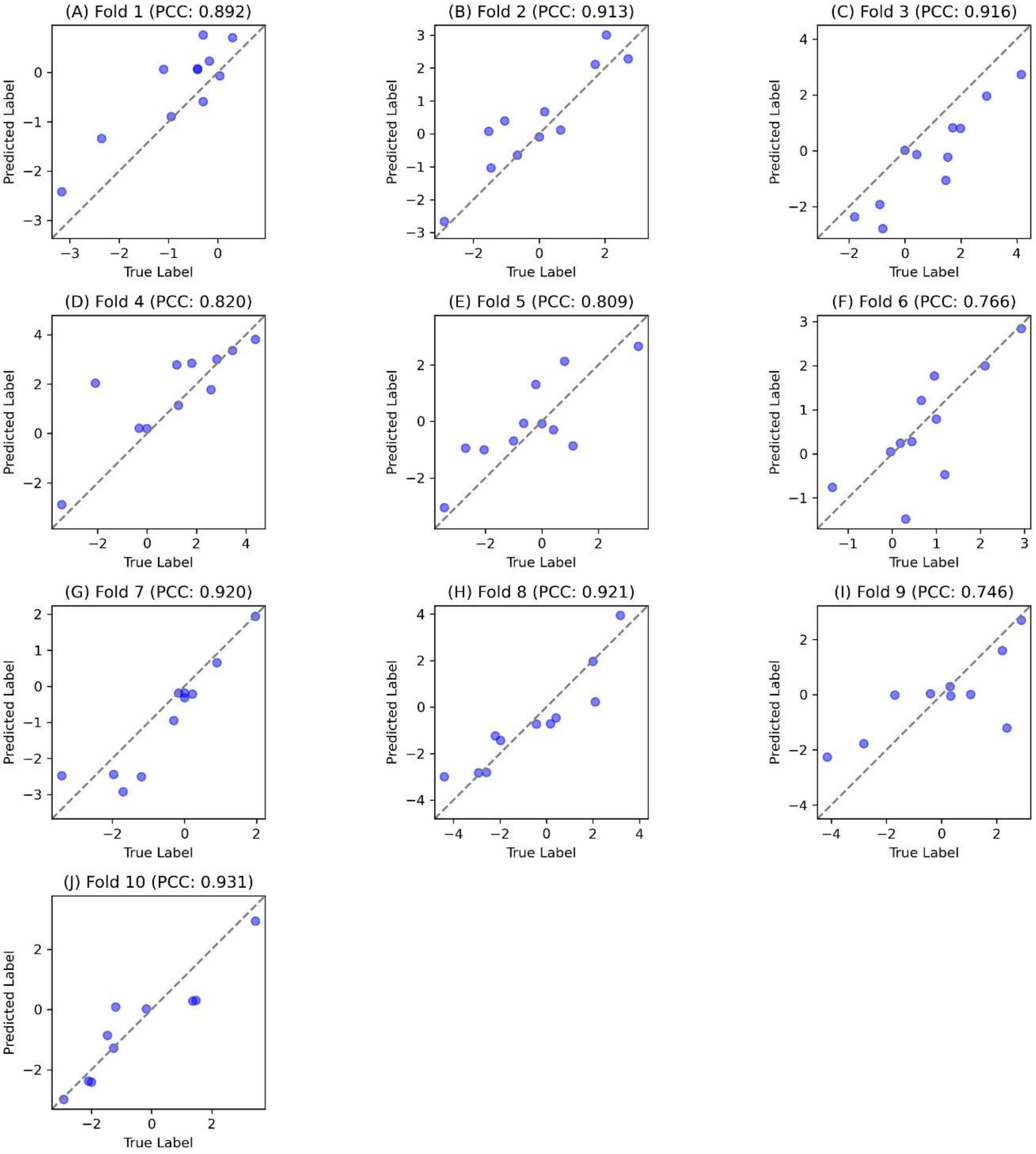


**Figure S3**. Scatter plots of predicted vs. true labels across ten-fold cross-validation on the SSDNA dataset. **(A)** Fold 1 (PCC: 0.892), **(B)** Fold 2 (PCC: 0.913), **(C)** Fold 3 (PCC: 0.916), **(D)** Fold 4 (PCC: 0.820), **(E)** Fold 5 (PCC: 0.809), **(F)** Fold 6 (PCC: 0.766), **(G)** Fold 7 (PCC: 0.920), **(H)** Fold 8 (PCC: 0.921), **(I)** Fold 9 (PCC: 0.746), **(J)** Fold 10 (PCC: 0.931).

Figure S3 provides a detailed assessment of a predictive model's performance on the SSDNA dataset using ten-fold cross-validation. Each subplot (A–J) displays a scatter plot of predicted versus true labels for a specific fold, with the Pearson Correlation Coefficient (PCC) ranging from 0.746 (Fold 9) to 0.931 (Fold 10). The x-axis represents true labels, and the y-axis denotes predicted labels, both spanning approximately -4 to 4. A dashed line (y=x) marks perfect prediction. Folds 7, 8, and 10 exhibit the highest PCC values (≥0.920), with data points closely aligned along the diagonal, indicating robust predictive accuracy. In contrast, Folds 6 and 9 show lower PCC values (0.766 and 0.746), reflecting increased dispersion and potential limitations in model generalization, possibly due to data variability or outliers. The predominantly high PCC values across most folds demonstrate the model's reliability, while the observed variability emphasizes the critical role of cross-validation in evaluating performance consistency. These findings highlight areas for potential model refinement to enhance predictive stability across diverse data subsets.

### Supplementary Figure S4.

#### Figure S4. Performance metrics across 10-fold cross-validation on the SSDNA dataset.

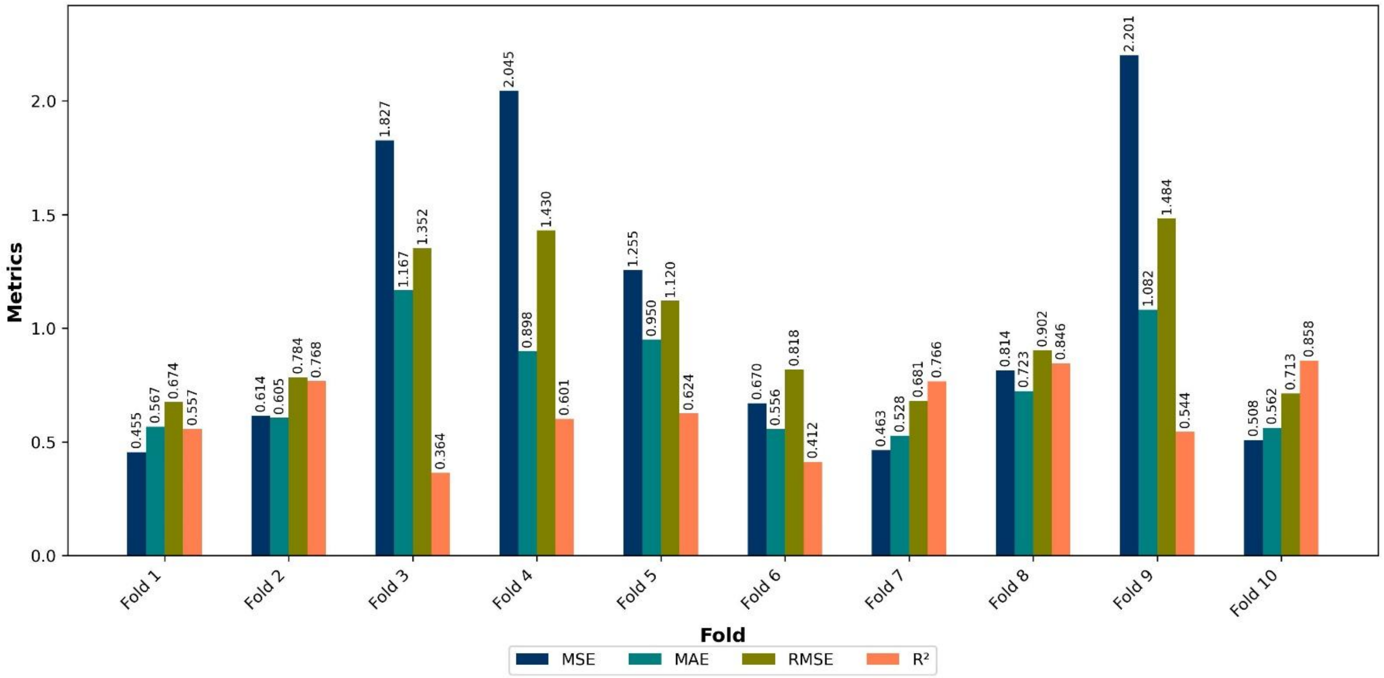


**Figure S4**. Performance metrics across 10-fold cross-validation on the SSDNA dataset.

Figure S4 presents a bar chart evaluating a predictive model's performance on the SSDNA dataset across ten-fold cross-validation, using four metrics: MSE (dark blue), MAE (teal), RMSE (olive), and $R^2$ (coral). Each fold (1–10) is plotted on the x-axis, with the y-axis scaled from 0 to 2.5 to capture metric variability. MSE ranges from 0.463 (Fold 7) to 2.201 (Fold 9), indicating significant error variation, while MAE (0.528–1.167) and RMSE (0.674–1.484) further highlight predictive inconsistency, peaking in Fold 9. $R^2$ values span 0.364 (Fold 3) to 0.858 (Fold 10), reflecting variable explanatory power, with Fold 10 demonstrating the best fit. The Pearson Correlation Coefficient (PCC, 0.746–0.931, not shown) and p-values ($\leq$0.013) from SSDNA108.xlsx confirm statistical robustness. Fold 9's elevated errors suggest potential data outliers, while Fold 10's metrics indicate optimal performance. This analysis underscores the model's reliability, with variability across folds warranting further investigation for refinement.
